# Integrated Multi-Omics Data Reveals the Molecular Subtypes of Prostate Cancer

**DOI:** 10.1101/2021.05.31.446411

**Authors:** Jialin Meng, Xiaofan Lu, Chen Jin, Yujie Zhou, Qintao Ge, Meng Zhang, Jun Zhou, Zongyao Hao, Fangrong Yan, Chaozhao Liang

## Abstract

Prostate cancer (PCa), the second most common male malignancy, is the fifth leading cause of cancer-related death and places notable burdens on medical resources. Most of the previous subtypes only focused on one or fewer types of data or ignored the genomic heterogeneity among PCa patients with diverse genetic backgrounds. Therefore, it is essential to precisely identify the specific molecular features and judge potential clinical outcomes based on multi-omics data. In the current study, we first identified the PCa multi-omics classification (PMOC) system based on the multi-omics, including mRNA, miRNA, lncRNA, DNA methylation, and gene mutation, using a total of ten state-of-the-art clustering algorithms. The PMOC1 subtype, also called the inflammatory subtype, contains the highest expression levels of immune checkpoint proteins, moderate activated immune-associated pathways. The PMOC2 tumor-activated subtype demonstrated the worst prognosis, which might be impacted by the activated cell cycle and DNA repair pathways, and also characterized by the most genetic alterations of mutant TP53, mutant APC and copy number alteration of 8q24.21 region. The PMOC3 subtype is likely to be a balance subtype, with the activated oncogenic signaling pathways, including hypoxia, angiogenesis, epithelial mesenchymal transition, and PI3K/AKT pathways. As well as with the activated proinflammatory pathways, including IL6/JAK/STAT3, IL2/STAT5, Notch and TNF-α signaling. Additionally, PMOC3 subtype also linked with the activation of the androgen response and the high response rate of ARSI treatment. Taken together, we defined the PMOC system for PCa patients via multi-omics data and consensus results of ten algorithms, this multi-omics consensus PCa molecular classification can further assist in the precise clinical treatment and development of targeted therapy.

## Introduction

Prostate cancer (PCa), the second most common male malignancy, is the fifth leading cause of cancer-related death among men and places notable burdens on medical resources. According to the latest data from the World Cancer Research Fund, approximately 15% of new male cancer cases are PCa, and there are 1.3 million newly diagnosed PCa cases per year. Moreover, the incidence and mortality of PCa vary significantly between regions and countries: the highest mortality is in Guadeloupe, with an age-standardized rate of 189.1/10^5^ ^1^, and the incidence and mortality in most developed regions are higher than those in other countries ^2^. The incidence rate of PCa increases with aging, and nearly 97% of new cases are diagnosed in humans older than 50 ^1^. However, high incidence and mortality may be a result of overdiagnosis with increasing prostate-specific antigen (PSA) testing, especially in developed countries^3^. Apart from these, race differences are an important feature of PCa incidence: African Americans are more likely to develop PCa at any age than other racial groups ^4^ and suffer from a higher risk of cancer-specific death^5^. In China, almost 7.1/10^5^ populations suffer from the burden generated by PCa. In addition, the incidence rate in urban areas (10.06/10^5^) is higher than that in rural areas (4.79/10^5^) ^6, 7^.

The clinical outcome and behavior of PCa are variable. Some localized PCa progresses to invasive cancer and ultimately leads to death, while other types of PCa remain indolent and can be cured by drugs or surgery. Therefore, it is important to distinguish different risk-level patients with PCa. There have been several risk classification systems based on clinical information and pathological parameters, including PSA and Gleason score^8, 9^. The molecular subtype is increasingly studied to divide different types of cancers into different subgroups, and huge progress has been made in this field. For example, Hedegaard et al. ^10^ classified non-muscle invasive bladder cancer into three molecular subtypes, and one of them was validated to be related to good prognosis. Human epidermal growth factor receptor 2 (HER2) is a signature molecule for screening suitable patients with breast cancer for chemotherapy ^11^. With the comprehensive utilization of HER2 in the clinic, disease-free and overall survival rates have been greatly prolonged ^12^. Therefore, it is essential to precisely identify the specific molecular features and judge potential clinical outcomes to help select appropriate treatment strategies for PCa patients. The characterization of PCa patients has been performed by various groups. You et al.^13^ described a prostate cancer classification system (PCS) defined by the activation signatures of 14 PCa-associated biological pathways. PCS1 tumors exhibited the worst prognosis, with resistance to androgen receptor signaling. PCS2 and PCS3 all have favorable clinical outcomes, but PCS3 has more bone metastasis. The Cancer Genome Atlas Research Network indicated a classifier of seven genetically distinct subtypes from 333 primary PCa patients, with differences in ERG fusion, ETV1/ETV4/FLI1 fusion, or SPOP, FOXA1, IDH1 mutations^14^, but still missed approximately 30% of PCa patients with undefined characteristics. They reported that AR activity varied widely, patients with SPOP and FOXA1 mutant tumors had the highest levels of AR-induced transcripts, and the actionable lesions of the PI3K/MAPK pathway or DNA repair pathway were approximately 25% and 19%, respectively. Zhao et al. ^15^ worked on the PAM50 classifier for prostate cancer classification and separated 495 PCa patients from the TCGA-PRAD cohort into luminal A, luminal B, and basal-like subgroups, revealing that the luminal B subtype suffers the worst recurrence-free survival and is significantly associated with the response to androgen deprivation therapy. However, most of the previous subtypes only focused on one or fewer types of data or ignored the genomic heterogeneity among PCa patients with diverse genetic backgrounds.

In the current study, we considered relapse-free associated features from the multiple omics data of the TCGA-PRAD cohort and constructed the PCa multi-omics classification (PMOC) system with diverse clinical outcomes and molecular characteristics with ten clustering algorithms.

## Methods and materials

### Obtaining multiple omics data

We used the TCGA-PRAD cohort as the training cohort, and data on mRNA expression, lncRNA expression, miRNA expression, DNA methylation and somatic mutations were enrolled to identify the different subtypes linked with relapse status and time. The row counts of mRNA and lncRNA were downloaded through the R package “*TCGAbiolinkS*” ^16^. For the mRNA expression component, the gene symbols were transformed after mapping the Ensembl IDs with the GENCODE27 annotation file. The lncRNAs were recognized via Vega (http://vega.archive.ensembl.org/) and recorded as noncoding, 3prime overlapping ncRNA, antisense RNA, lincRNA, sense intronic, sense overlapping, macro lncRNA, and bidirectional promoter lncRNA subtypes. The raw counts of mature miRNAs were downloaded from UCSC Xena and translated to mature miRNA names in miRbase version 21.0 via the R package “*miRNAmeConverter*”. The raw read counts were transferred into transcripts per kilobase million (TPM) values. The Illumina DNA methylation 450 data was also downloaded from UCSC Xena. Somatic mutations, clinicopathological features and relapse-free survival (RFS) data were downloaded from cBioPortal (https://www.cbioportal.org/). After merging the available samples in all omics and clinical features, a total of 486 PCa patients were left for subsequent analysis.

Four PCa patient cohorts recorded available clinicopathological information, and RFS data were also enrolled as the external validation cohort, including the GSE54460, GSE70770, MSKCC and GSE116918 cohorts (**Table S1**). The gene symbol for each cohort was transferred from the probe ID according to the corresponding annotation file from each platform. For one gene symbol that was mapped by multiple probes, the mean value was calculated as the expression level. The detailed clinical information of the TCGA-PRAD cohort and four external validation cohorts are listed in **Table 1**.

### Integration of multi-omics and recognition of subtypes

We performed the recognition of subtypes of PCa patients based on mRNA expression, lncRNA expression, miRNA expression, DNA methylation and somatic mutations data. The TPM expression data of mRNA, lncRNA and miRNA were first transformed by log2 calculation. For the DNA methylation data, we only focused on the probes located at the promoter region CpG islands. For the gene mutation matrix, we considered the gene as having mutant status if it contained any of the following nonsynonymous variations: frameshift deletion/insertion, in-frame deletion/insertion, missense/nonsense/nonstop mutation, splice site or translation start site mutation. To identify the subtypes that could mostly separate the different clinical outcomes for PCa patients, we selected the top factors most highly related to RFS, including 1526 mRNA, 242 lncRNA, 30 miRNA, and 1073 DNA CpG methylation sites, and 23 mutant genes had a mutation rate higher than 0.3. Subsequently, we selected the number of subtypes based on the prediction results of the clustering prediction index (CPI) ^17^ and Gaps-statistics ^18^. Then, we performed clustering via 10 state-of-the-art multi-omics clustering algorithms, including iClusterBayes, moCluster, CIMLR, IntNMF, ConsensusClustering, COCA, NEMO, PINSPlus, SNF, and LRA. The final subtype clustering combines the results of the 10 algorithms mentioned above to make the clustering more robust.

### Specific activated signaling pathways among subtypes

To reveal the specific characteristics of each subtype, we compared the distribution of oncogenic pathways, metabolic pathways and infiltration of immunocytes in the newly recognized subtypes. The enrichment score of single-sample gene set enrichment analysis (ssGSEA) for all concerned pathways was calculated by the R package “GSVA” ^19^. For the oncogenic pathways, we used the 50 tumor-associated gene sets from HALLMARK^20^. For the metabolic pathways, we enrolled a total of 113 gene sets from a published article^21^. We also assessed the infiltration rate of immunocytes with the CIBERSORT algorithm^22^, and the results of immunocytes with an average infiltration value of 0.01 were discarded. The functional proteomic data of 218 proteins were downloaded from The Cancer Proteome Atlas (TCPA) project, which used a reversed-phase protein array to detect the level of proteins combined with antibodies^23^. To compare the mRNA or protein levels between different subtypes, the average level of each subtype was first determined, and then the level of each gene or protein among the subtypes was transferred to the normal distribution to make all of the results comparable.

### Transcriptional regulatory networks among three PMOC subtypes

We assessed the activation of tumorigenesis pivotal transcriptional regulatory networks (regulons) composed of the transcription factor and its downstream induced or repressed target genes. The analysis of regulons was conducted by the package “*RTN*”, as previously reported ^24^. Moreover, we evaluated the impact of 71 chromatin remodeling-associated genes, which are regarded as potential gene regulators ^25^. Mutual information analysis and Spearman’s correlation analysis were applied to assess the potential associations among regulators and downstream target genes through the gene expression matrix, and associations identified by false discovery rate (FDR)-adjusted p-values higher than 0.00001 evaluated by permutation analysis were discarded. In addition, unstable associations were eliminated by bootstrap analysis (1,000 resamplings, consensus bootstrap >95%), and the weakest associations in triangles of two regulators and common target genes were removed by data processing inequality (DPI) filtering. Regulon activity scores for all samples were calculated by two-tailed GSEA.

### Evaluation of AR activity and response androgen deprivation (ADT) therapy

Two previously reported AR activity signatures were collected, and the activation score was also calculated by GSEA. Farzana et al.^26^ reported the AR Activity (AR-A) score, which is taken as a weighted linear sum of 9 canonical AR transcriptional target genes (KLK3, KLK2, FKBP5, STEAP1, STEAP2, PPAP2A, RAB3B, ACSL3, NKX3-1). Haley et al. refined the AR activation signature with 27 genes that show robust activation or inhibition of expression upon androgen stimulation, including KLK3, TMPRSS2, NKX3-1, KLK2, GNMT, PMEPA1, MPHOSPH9, ZBTB10, EAF2, CENPN, C1orf116, ACSL3, PTGER4, ABCC4, NNMT, ADAM7, FKBP5, ELL2, MED28, HERC3, MAF, TNK1, GLRA2, MAPRE2, PIP4K2B, MAN1A1, and CD200. A clinical cohort containing the gene matrix and response status to ADT therapy was also employed to evaluate the diverse response to ADT among three PMOC subtypes^27^. The estimated IC_50_ of bicalutamide, an AR signaling inhibitor (ARSI), in patients was calculated by the R package “*pRRophetic*” after comparison with the recorded response IC_50_ value of the drug in urinary tumor cell lines by ridge regression and 10-fold cross-validation prediction accuracy value ^28^.

### Characteristics of genetic alterations among subtypes

The diversity distribution of mutant genes among subtypes was compared by Chi-square test and displayed with a triangle plot, in which the coordinates of each point represent the mutation frequency of the gene, and the closer a point is to a corner of the triangle, the more mutation is present in the subtype. The size of the point is linked with the total mutation frequency of the gene. The mRNA expression among wild-type and mutant APC samples was compared by the Wilcoxon test, and the cumulative recurrent events were displayed by the R package “*Survival*”. The copy number segment data were downloaded from FireBrowse (http://firebrowse.org/) and displayed by the R package “maftools”.

### Statistical analyses

All statistical tests were executed by R (Version 4.0.2), including Fisher’s exact test for categorical data, a two-sample Mann-Whitney test for continuous data, a log-rank test with a Kaplan-Meier curve, and Cox proportional hazards regression for hazard ratio (HR) with a 95% confidence interval (95% CI). The biological differences in different LumB subtypes identified were evaluated by GSEA through the R package “*clusterProfiler*” ^29^. Specifically, GSEA was performed based on a pre-ranked gene list sorted by log2FoldChange derived from limma differential expression analysis. For the subtype identified in the external validation cohort, the top 300 specific marker genes for each subtype were used to separate the external cohort into subtypes by nearest template prediction (NTP) analysis^30^. Most of the above analytic processes are embedded in the R package “*MOVICS*”, which we recently developed for multi-omics integration and visualization^31^. For unadjusted comparisons, a two-sided test with *P* < 0.05 was considered to be significant.

## Results

### Identify the RFS relevant PCa subtypes

With the help of CPI and Gaps-statistics analysis, as well as the predefined PAM50 cluster for PCa patients, we found that when the number of clusters was three, the score from the two methods was more approximate (**Figure 1A**). Then, we preset the number of three for ten multi-omics integrative clustering analyses (**Figure 1B**) and further combined the clustering results of the ten algorithms via ensemble consensus for the robust subtypes (**Figure 1C**). Finally, we recognized three PCa subtypes with diverse clinical RFS outcomes, named PMOC1, PMOC2 and PMOC3 (**Figure 1D**). Patients in the PMOC2 subtype had the worst clinical outcome, while patients in the PMOC3 subtype had the longest recurrence-free survival time (PMOC1 vs. PMOC2, *P* < 0.001, PMOC1 vs. PMOC3, *P* < 0.001, PMOC2 vs. PMOC3, *P* < 0.001, **Figure 1D**). The distribution of the enrolled multi-omics data for subtype clustering is shown in **Figure 2**. The top ten diversity factors for each omics approach are listed on the right. We also compared the clinical features of the three subtypes. Patients in the PMOC2 subtype mostly had a higher Gleason score (9+10) than those in the PMOC1 and PMOC3 subtypes (61.8% vs. 23.0% vs. 9.7%, *P* = 0.012, **Table 2**), as well as a higher proportion of advanced pathology T stage patients (86.8% vs. 54.9% vs. 52.6%, *P* < 0.001 **Table 2**).

**Figure 1.**
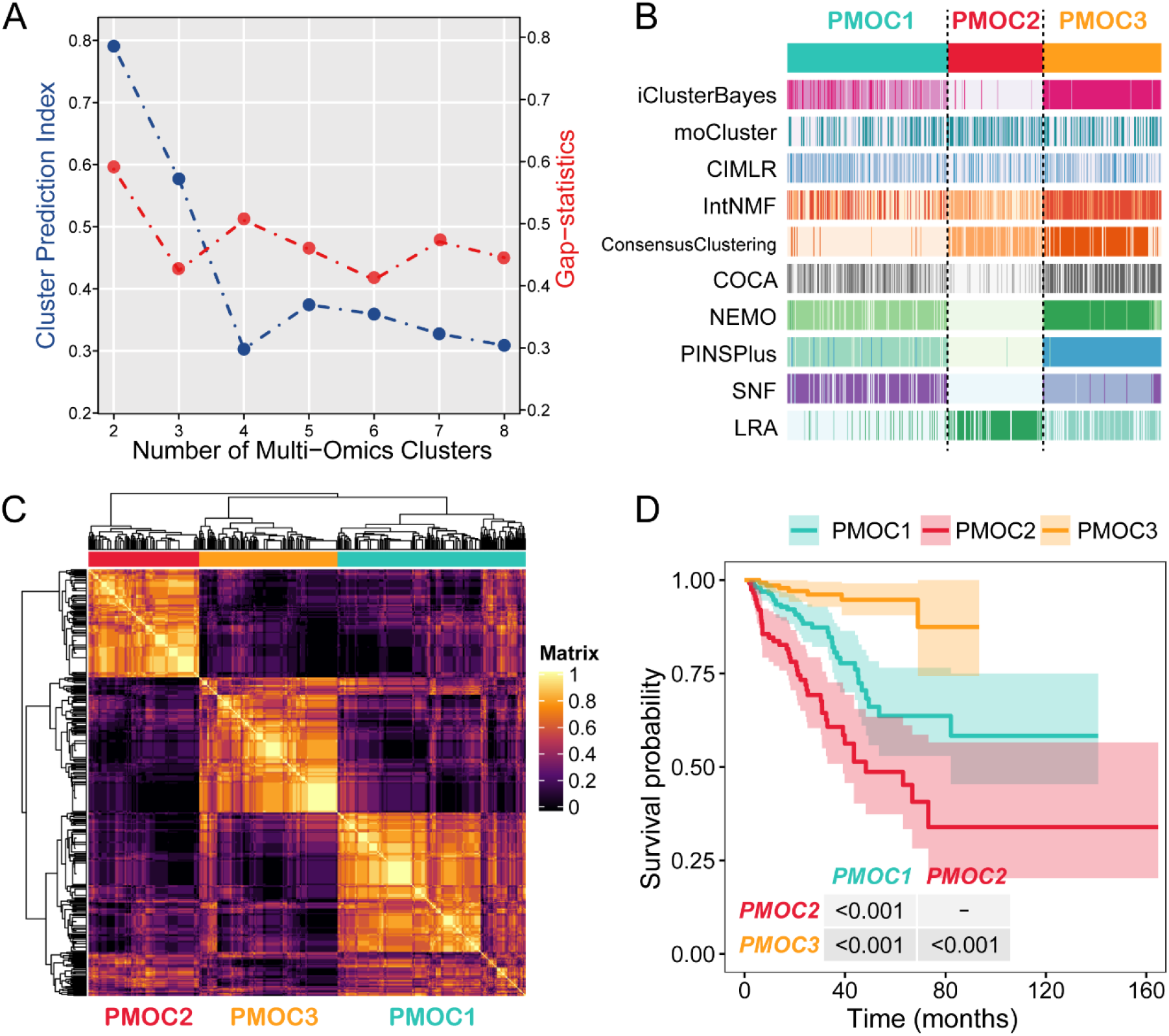
Recognition of the genomic subtypes in the TCGA-PRAD cohort. (A) Speculation regarding the appropriate clustering number by clustering prediction index and Gaps-statistics; (B) clustering of PCa patients via 10 state-of-the-art multi-omics algorithms; (C) consensus matrix for three clusters based on the 10 algorithms; (D) differential recurrence-free survival outcome in three subtypes evaluated by Kaplan-Meier survival analysis and the log-rank test.

**Figure 2.**
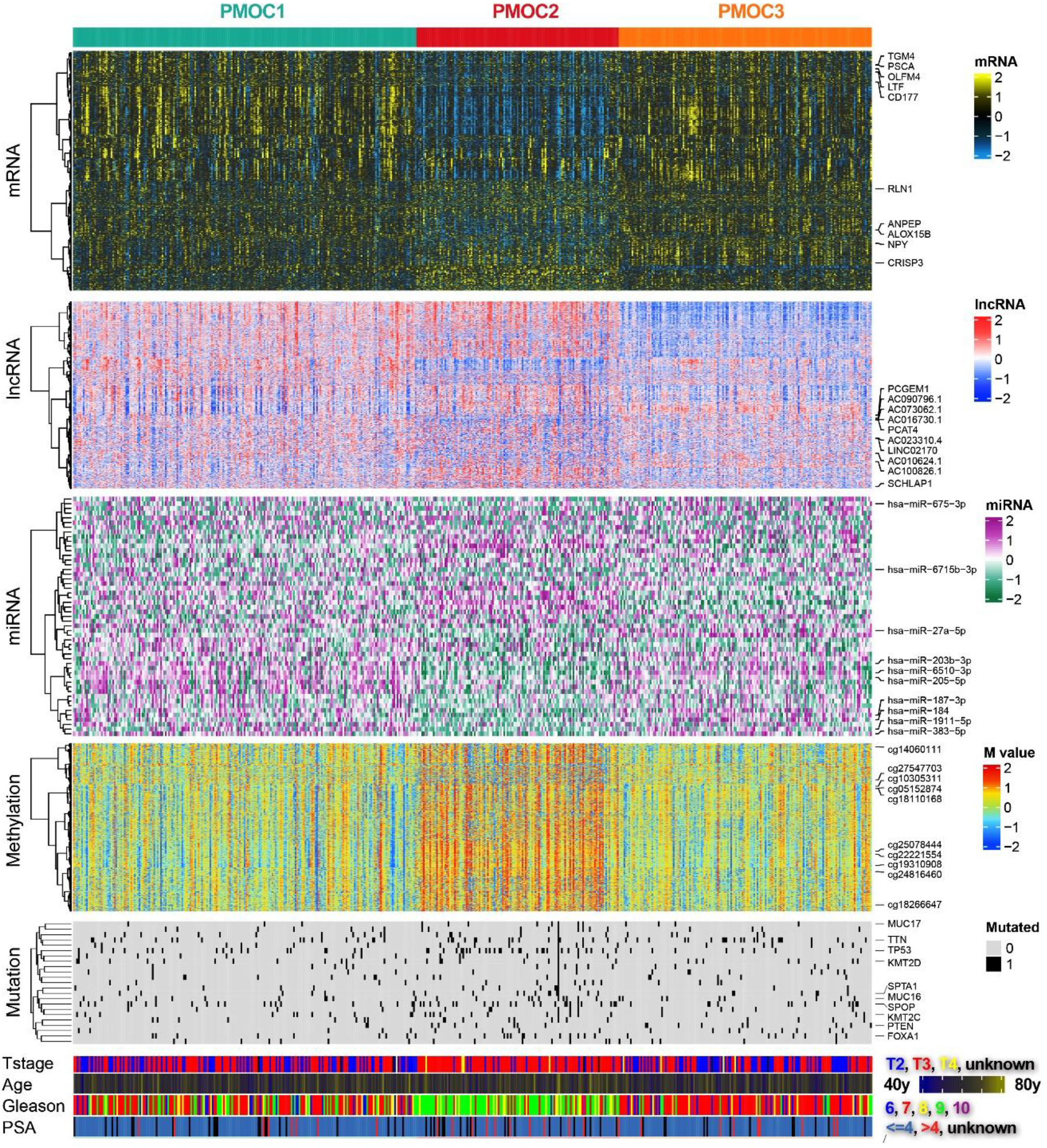
Integration of genomic subtype classifications from multi-omics data. Omics data of 1526 mRNAs, 242 lncRNAs, 30 miRNAs, 1073 DNA CpG methylation sites, and 23 mutant genes were considered for classification.

### Activated cell cycle-associated pathway foreboded the poor prognosis PMOC2 subtype

We first compared the activation status of 50 tumorigenesis-associated pathways in the three subtypes. We observed the significance of the G2M checkpoint, E2F target pathways in the PMOC2 subtype, and the two MYC target pathways and DNA repair pathway (**Figure 3A**). Activated cell cycle-associated pathways were positively linked with poor prognosis. We illustrated the G2M regulatory pathways in detail and revealed that the phosphorylated protein levels of p-CHK1 and p-CHK2 decreased in PMOC2, which weakened the inhibitory function of CDC25 components. Then, the mRNA, protein and phosphorylated protein levels of CDK1, a highly conserved serine/threonine kinase that plays a key role in cell cycle regulation, were increased (**Figure 3B**). For PMOC1, we observed that only a few pathways were activated, such as TNF-α signaling, IL6 JAK STAT3 signaling and IL2 STAT5 signaling. These results might reflect that PMOC1 is linked to the immune response. Although the PMOC3 subtype also presented activation of the immune-associated pathway, we also observed high activation of several oncogenic pathways, including the PI3K/AKT, hypoxia, apoptosis, angiogenesis, EMT, P53, hedgehog and Wnt β-catenin signaling pathways. Combined with the best prognosis of the PMOC3 subtype, there might be a sort of “balance” in the tumor environment to combat tumor cells, leading to a good clinical outcome in PMOC3. For PI3K/AKT signaling, we observed higher mRNA levels of PI3K components in PMOC3 and further promoted the mRNA level of AKT components, as well as the protein levels of AKT and phosphorylated AKT. The phosphorylated mTOR and mTOR levels are also activated and ultimately lead to the promotion of cell growth (**Figure 3C**).

**Figure 3.**
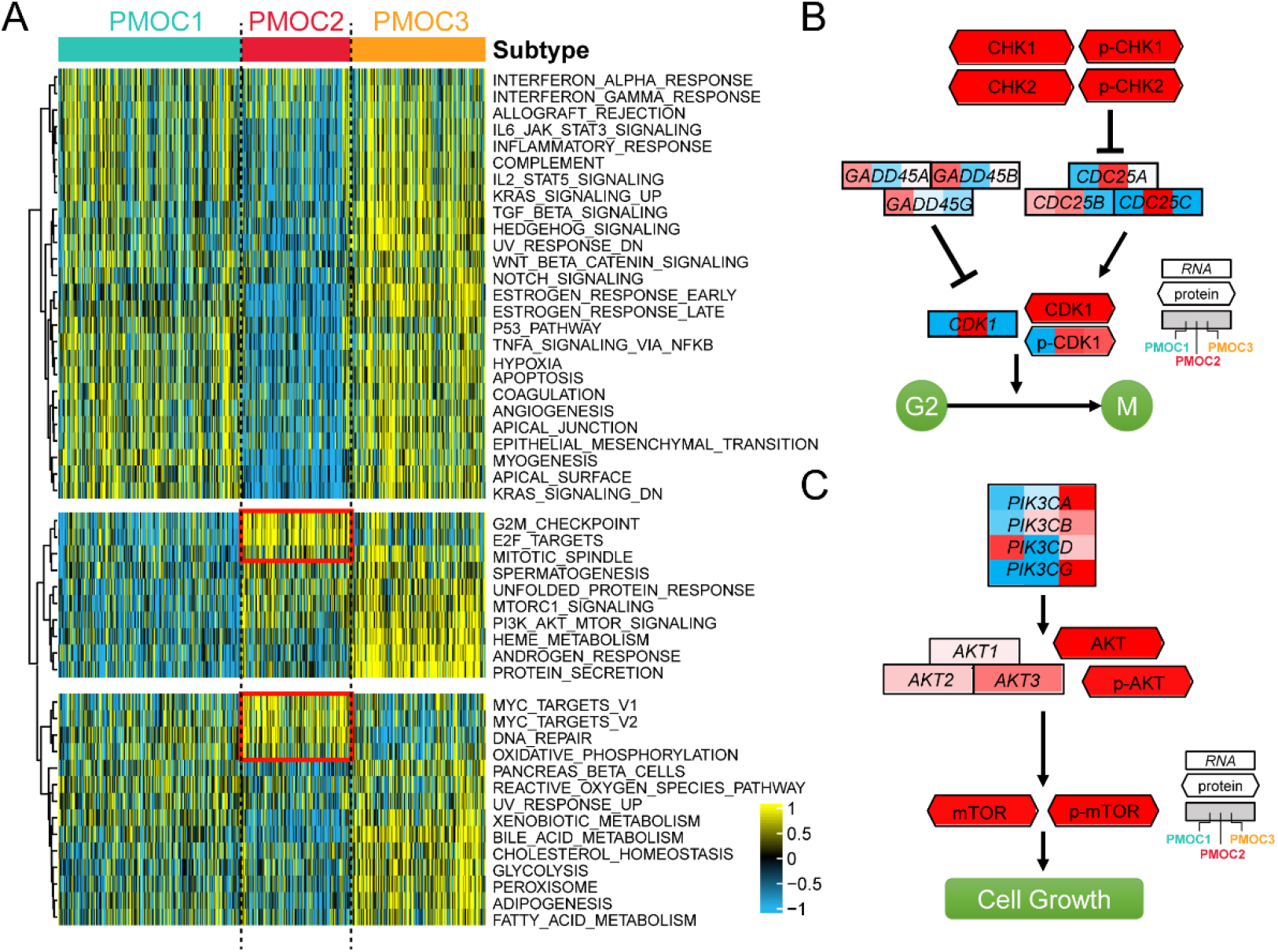
Differential activity of tumor-associated pathways across PCa genomic subtypes. (A) Heatmap of 50 differentially activated HALLMARKER pathways; (B) G2M pathways activated in the PMOC2 subtype at both the mRNA and protein levels; (C) PI3K/AKT pathways activated in the PMOC3 subtype at both the mRNA and protein levels.

### Diversity metabolism pathways and immunocyte infiltration represent quite different tumor microenvironments

To further characterize the newly defined PCa subtypes, we compared the activation status of metabolic pathways. Interestingly, we revealed that the nicotinamide adenine dinucleotide (NAD) biosynthesis and cyclooxygenase (COX) arachidonic acid (ARA) metabolism pathways were activated in the PMOC1 subtype (**Figure 4A**). For the PMOC2 subtype, pathways associated with cell cycle activation were consistent with the findings in hallmark tumor pathways, including pyrimidine metabolism and biosynthesis, biotin metabolism, and oxidative phosphorylation (**Figure 4A**). In the PMOC3 subtype, glycogen metabolism and amino acid metabolism-associated pathways were highly activated, which also reflected the numerous and vibrant tumor pathways in the tumor microenvironment revealed by the hallmark results (**Figure 4A**).

**Figure 4.**
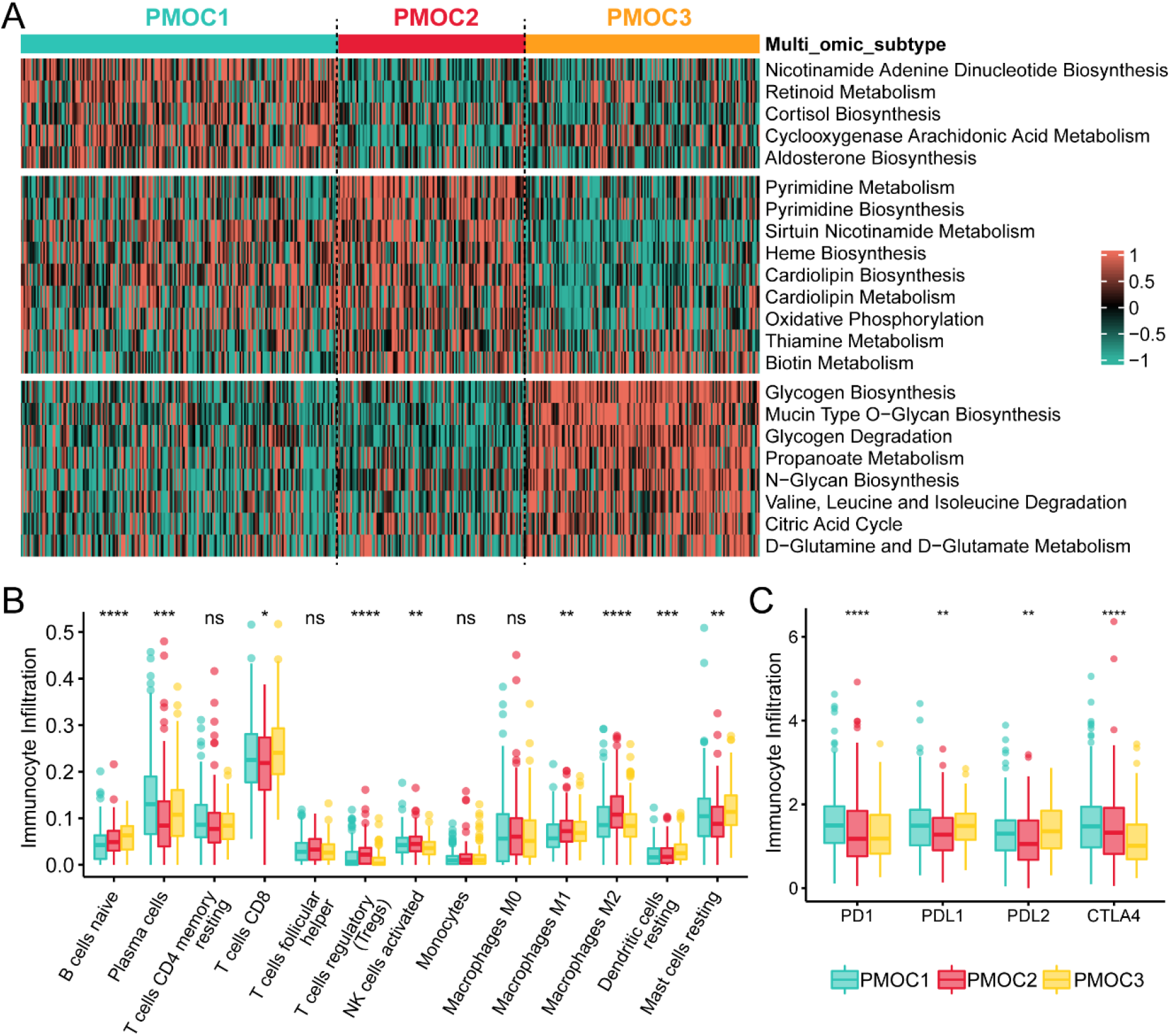
Special metabolism and immune characteristics across PCa genomic subtypes. (A) Heatmap of subtype-specific metabolism signaling pathways; (B) Differential infiltration of 13 immunocytes among three subtypes; (C) Expression patterns of four immune checkpoints across three PCa subtypes. ns, not significant; *, *P* < 0.05; **, *P* < 0.01; ***, *P* < 0.001; ****, *P* < 0.0001.

In addition, we compared the infiltration of immunocytes and the expression of immune checkpoint differences. We observed that although the PMOC3 subtype had higher infiltration of resting CD4+ memory T cells, naïve B cells, M1 macrophages, resting dendritic cells, and resting mast cells, the PMOC1 subtype seemed to exhibit more highly activated immune infiltration, represented by higher infiltration of plasma cells, CD8+ T cells, and activated NK cells (**Figure 4B**), as well as higher expression of PD1, PDL1 and CTLA4 (**Figure 4C**). Anti-inflammatory immunocytes highly infiltrated PMOC2 subtypes, including Tregs, follicular helper T cells, and M2 macrophages.

### Genetic alteration enriched the phenotype of three subtypes

Genetic alteration notably impacts the shaping of subtypes. As shown in **Figure 2**, we observed the enrichment of patients with gene mutations in the PMOC2 subtype, and the total tumor mutant burden was highest in the PMOC2 subtype (*P* < 0001, **Figure S1**). In detail, we listed a total of 13 genes with a significantly different distribution of wild-type or mutant genes in the three subtypes (**Table S2**). A total of 56 patients had TP53 mutations (11.5%), and the PMOC2 subtype contained most patients with this mutation (PMOC2: 23.6%, PMOC1: 8.6%, PMOC3: 5.8%, *P* < 0.001). Mutant SPOP also mostly presented in the PMOC2 subtype (PMOC2: 18.7%, PMOC1: 9.6%, PMOC3: 7.8%, *P* = 0.0138). We further focused on mutations in the APC gene, and 7 of 10 mutant patients presented with the PMOC2 subtype (**Figure 5A**). The APC protein is an antagonist of the Wnt signaling pathway and acts as a tumor suppressor^32^. Mutant APC resulted in lower expression (*P* = 0.0026, **Figure 5B**) and was associated with a poor prognosis (HR: 3.61, 95% CI: 1.32-9.92, *P* = 0.013, **Figure 5C**).

**Figure 5.**
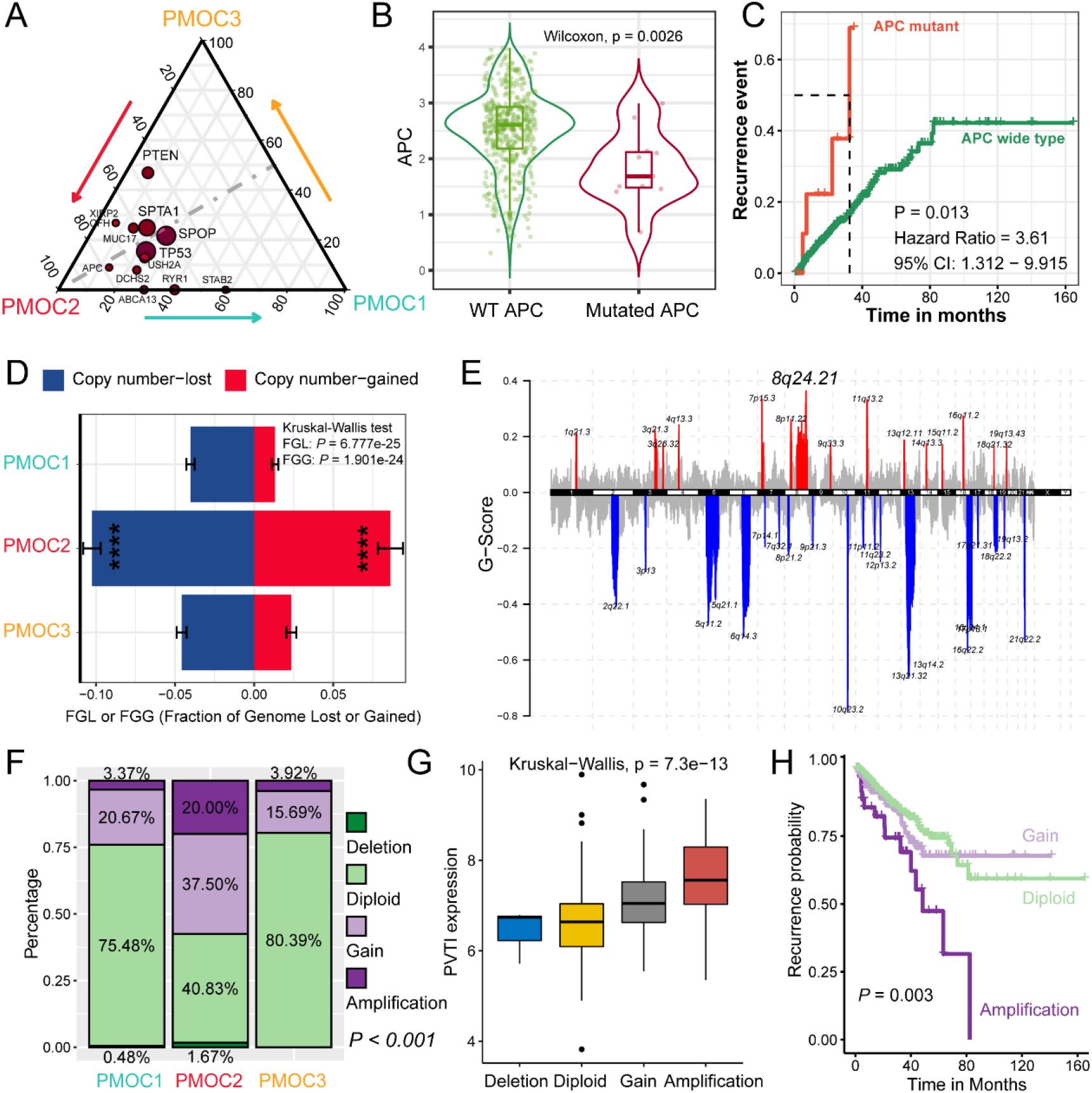
Genetic alteration landscape among PCa genomic subtypes. (A) Triangle plot showing the differential mutation frequency of tumor-driven mutant genes; (B) Mutated APC caused the downregulation of self mRNA expression; (C) Mutated APC resulted in the greater accumulation of recurrence events; (D) Disparities in copy numbers gained and lost in PMOC subtypes; (E) Plot of G scores calculated by GISTIC 2.0 for genomic regions amplified (red) or deleted (blue) in the PMOC2 subtype, and G scores were defined as the amplitude of copy number multiplied by frequency across patients; (F) Copy number alteration of PVT1 in three PMOC subtypes; (G) Correlation of copy number alteration with mRNA expression of PVTI; (H) Amplification of PVTI indicated the worst clinical outcome.

We also evaluated the copy number alteration in the three subtypes and revealed that both the lost and gained copy numbers were significantly increased in the PMOC2 subtype compared with the PMOC1 and PMOC3 subtypes (**Figure 5D**). Interestingly, we observed that the gained copy number in PMOC2 was mostly located at the 8q24.21 region but was not observed in the same region in patients belonging to the PMOC1 or PMOC3 subtype (**Figure 5E, Figure S2**). PVT1 is an important gene located in the 8q24.21 region, and its expression in tumor tissues is significantly higher than that in normal tissues across cancers (**Figure S3**). We found that the amplification and gain alteration of PVT1 occurred mostly in PMOC2 subtype patients (*P* < 0.001, **Figure 5F**), the mRNA expression level of PVT1 was positively correlated with the change in copy number (*P* < 0.001, **Figure 5G**), and patients with the amplification expression of PVT1 suffered from the worst recurrence-free survival time (*P* = 0.003, **Figure 5H**).

### PMOC3 subtype can benefit more from ADT therapy

Transcript factors always play important roles in the genesis and progression of tumors. Therefore, we evaluated the activation of 23 regulons and 71 chromatin remodeling potential regulons. We observed a quiet difference in activated regulons in different PMOC subtypes. Patients with PMOC2 are likely regulated by the human Fox gene family (FOXM1, FOXA1), as well as the chromatin remodeling-linked genes KAT2A, SMYD3, EZH2 and NSD2. For patients in the PMOC3 group, we were first attracted by the activated regulation of AR, EGFR, and HIF1A (**Figure 6A**). Based on the GSEA calculation, we observed significantly enriched AR activation in PMOC3 via the ARA score (*P* < 0.001, **Figure 6B**) and AR activation signature (*P* < 0.001, **Figure 6C**). For the ADT-treated Abida cohort, the NTP algorithm was used to represent the three PMOC subtypes with 300 subtype-specific genes. Interestingly, we observed similar activated pathways as those in the TCGA cohort. PMOC2 demonstrated the activation of cell cycle-associated pathways, and PMOC3 was linked with the activated androgen, estrogen and PI3K/AKT signaling pathways (**Figure 6D**). We also revealed that patients in PMOC3 were more likely to respond to ADT therapy (response rate of 26.7% in PMOC3, **Figure 6E**), including bicalutamide (*P* = 0.089, **Figure 6F**).

**Figure 6.**
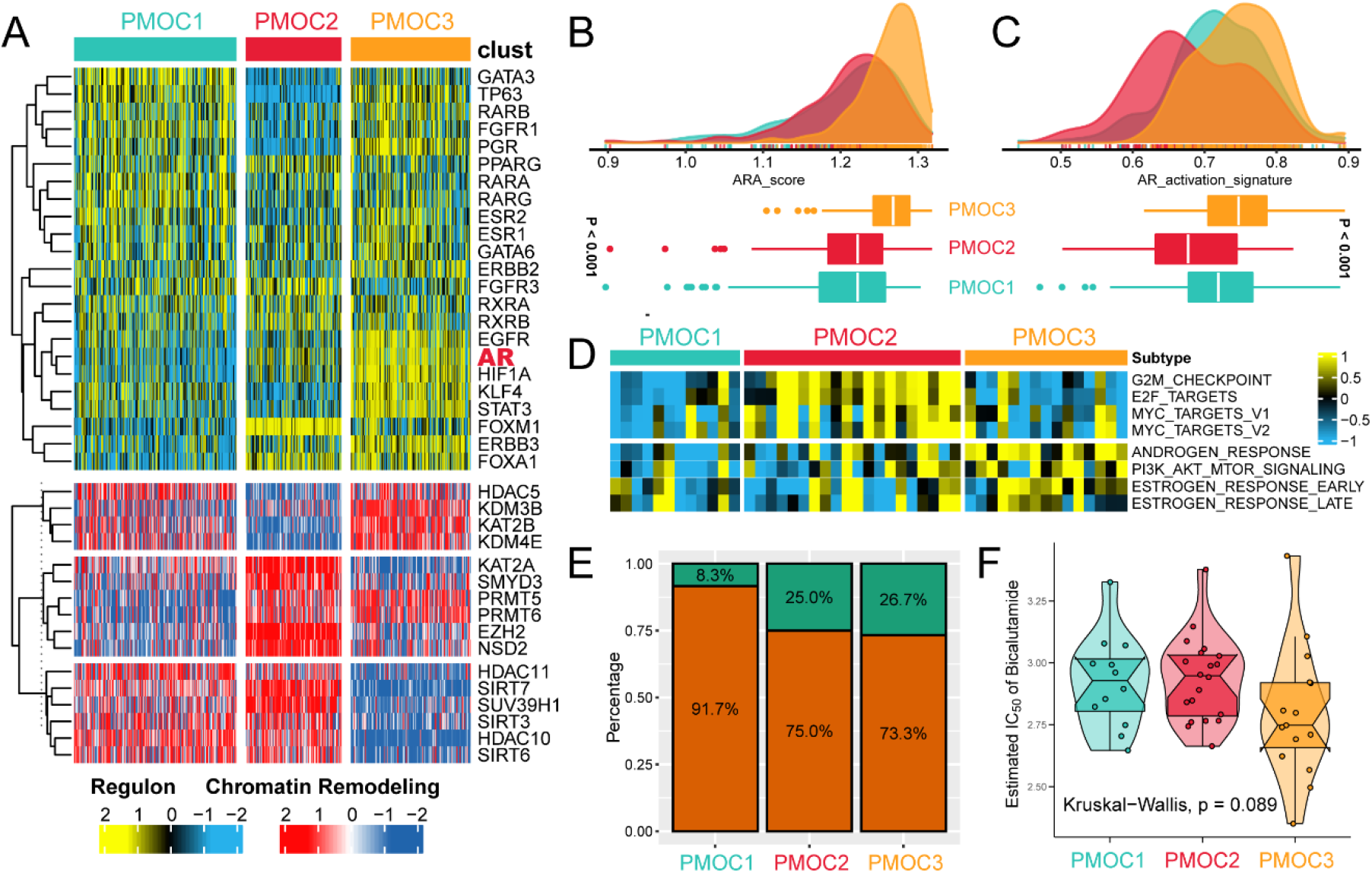
Differential response to ADT therapy across PCa genomic subtypes. (A) Regulon activity profiles of 23 transcription factors and potential regulators associated with chromatin remodeling. Activation of the AR signaling pathway evaluated by ARA score (B) and AR activation signature (C) among three PMOC subtypes in TCGA-PRAD cohort; (D) Differentially activated specific pathways in PMOC2 and PMOC3 validated in patients in the Abida cohort; (E) Patients in the PMOC3 subtype benefit more from ADT therapy; (F) Patients in the PMOC3 subtype are more suitable for bicalutamide treatment.

### Redefined subtypes and comparison with existing molecular classifiers

Based on the findings of the three subtypes, we defined the PMOC1 subtype as the tumor-inflammatory subtype because it has the moderate activation of the immune microenvironment, high infiltration of immunocytes and high expression of PD-1, PD-L1 and CTLA4. The PMOC1 subtype is tightly linked with the activated immune subgroup we previously identified, which has an activated immune environment and good prognosis^33^ (**Figure 7A**).

**Figure 7.**
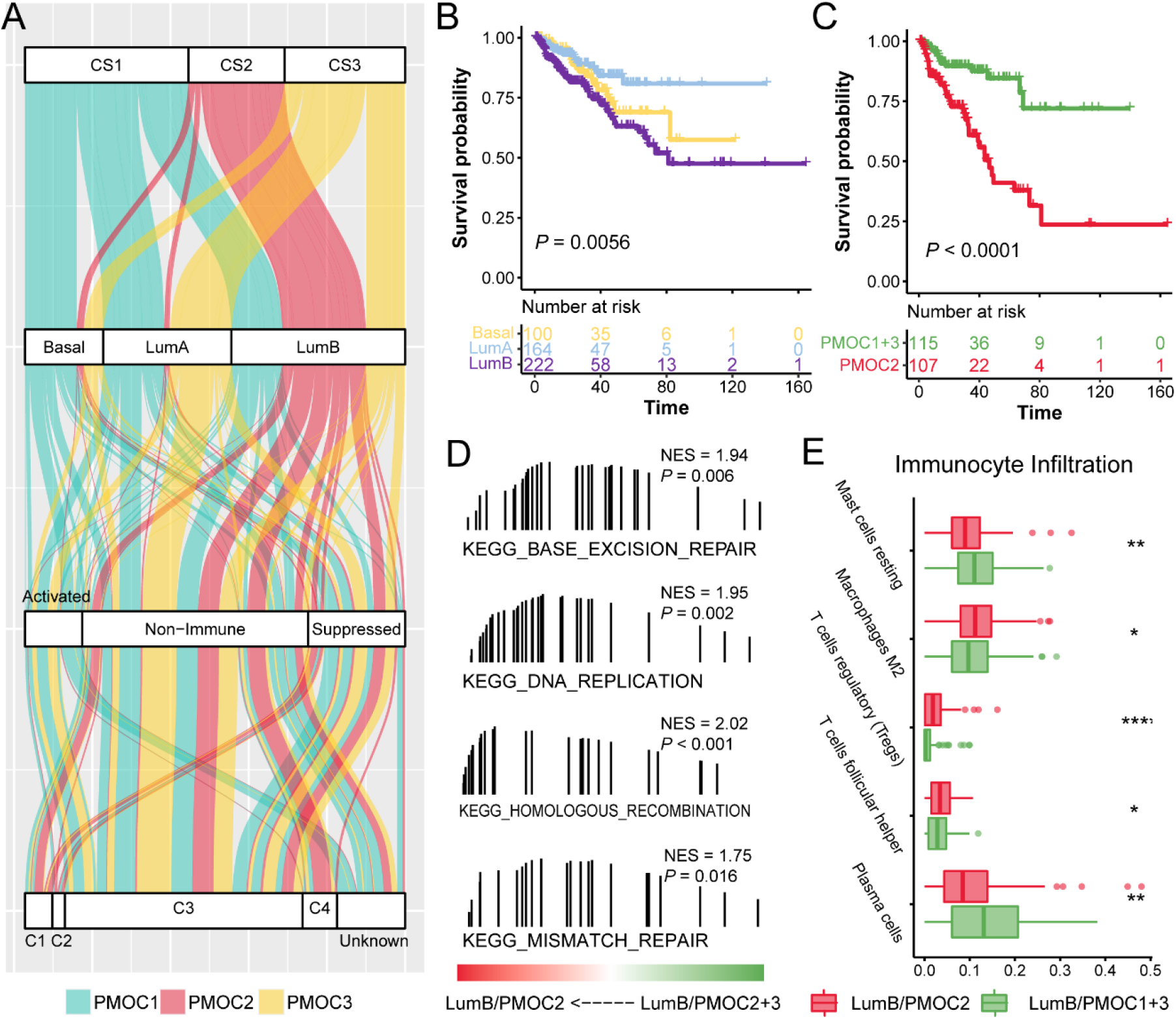
PMOC subtypes separated the PCa luminal B molecular subtype with diverse recurrence outcomes. (A) Overlap between PMOC subtypes, PAM50, immune activation subtypes and six immune molecular features. (B) Differential recurrence-free survival outcomes in the basal, LumA and LumB subtypes; (C) Differential recurrence-free survival outcome in the LumB/PMOC1+3 and LumB/PCS2 subtypes; (D) DNA replication and repair pathways activated in LumB/PMOC2 compared with LumB/PMOC1+3; (E) Differential infiltration of immunocytes in two separate LumB subtypes.

The PMOC2 subtype, named the tumor-activated subtype, demonstrates the activation of cell cycle-associated signaling pathways and dramatically alters TP53 mutation, SPOP mutation, APC mutation, and the 8q24.21 copy number. Then, its tumor-activated status leads to a poor prognosis.

The PMOC3 subtype involves the activation of numerous signaling pathways, including both proinflammatory and oncogenic pathways, as well as vibrant glycogen and amino acid metabolism. Multiangle activation results in a sense of “balance” in the tumor microenvironment and leads to a favorable clinical outcome. Therefore, the PMOC3 subtype was named the tumor-balance subtype.

We also compared the three PCa subtypes with PAM50 subtypes. The LumB subtype had the worst prognosis compared with the LumA and basal subtypes (*P* = 0.0056, **Figure 7B**). We found that the PMOC2 tumor-activated subtype mostly belongs to LumB and demonstrates the worst prognosis compared to the PMOC1 and PMOC3 subtypes, which means that the PMOC2 subtype can help recognize a subgroup of LumB patients with unfavorable clinical outcomes. As we speculated, LumB/PMOC2 patients had an unfavorable outcome compared with LumB/PMOC1+PMOC3 patients (*P* < 0.001, **Figure 7C**), and this difference might be impacted by the DNA repair and replication pathways (all *P* < 0.05, **Figure 7D**). We also found that LumB/PMOC2 cells had higher infiltration of anti-inflammatory immunocytes, M2 macrophages, Tregs and follicular helper T cells (**Figure 7E**).

### Validation of diverse clinical outcomes of subtypes in external cohorts

To reproduce the three subtypes, we selected the top 100 specific genes of each subtype that had the highest values compared with the other two subtypes (**Table S3**) and represented the separation of the inflammatory, activated and balanced subtypes (**Figure S4**). In the GSE54460 cohort, 42 patients belonged to the PMOC1 subtype, with moderate RFS; 30 patients were separated into PMOC2 subtypes and presented a poor prognosis, while PMOC3 subtype patients had the most favorable clinical outcome (*P* < 0.001, **Figure 8A**). We also compared clinical outcomes in the GSE70770, MSKCC and GSE116918 cohorts. Although the survival times of the PMOC1 and PMOC3 subtypes were comparable to each other, patients with the PMOC2 subtype had the worst clinical outcomes (all *P* < 0.05, **Figure 8**).

**Figure 8.**
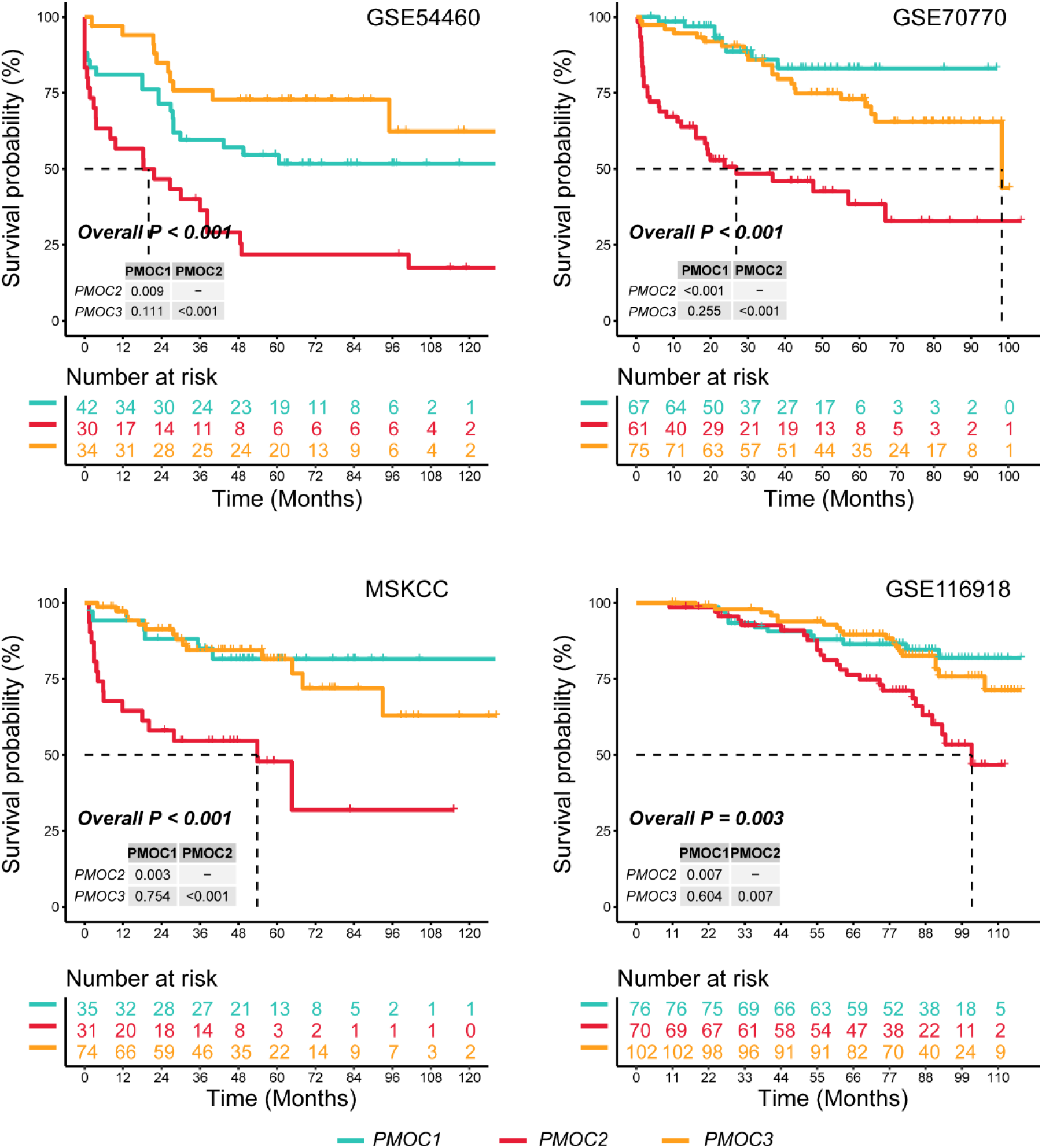
Validation of the 300-gene signature to reproduce the three PMOC subtypes in external cohorts. Kaplan-Meier curves and log-rank tests were used to display and compare recurrence-free survival in the three PMOC subtypes. The *P*-value between two subtypes was adjusted by the Benjamini and Hochberg step-up procedure.

We also conducted multiple Cox regression analysis to determine the prognostic value of PMOC subtypes after adjusting for the dominant clinical features. As shown in **Table 3**, we observed that patients in the PMOC2 group had the worst clinical outcomes, even after removing the potential bias caused by other factors in all five PCa patient cohorts (all *P* < 0.05).

## Discussion

PCa health burden is a serious problem worldwide, with a high recurrence rate and long-term discomfort for male patients. Physical fatigue, pain, emotional fatigue and depressive symptoms are excruciating for long-term PCa survivors and are impacted by clinical treatment and comorbidities^34^. Additionally, the bone metastases of advanced stage PCa patients can lead to severe pain, nerve compression and even anemia^35^.

In the current study, we first identified the PMOC system based on the multi-omics, including mRNA, miRNA, lncRNA, DNA methylation, and gene mutation, using a total of ten state-of-the-art clustering algorithms. Single omics data has been widely used for PCa classification and prognosis prediction, providing mild value in clinical research and treatment. In our prior study, we revealed that the infiltration of immunocytes, especially M2 macrophages, is linked with unfavorable prognosis in PCa patients^36^, and further study revealed the heterogeneous immune microenvironment in PCa patients. Patients with highly infiltrated immunocytes were separated into immune-activated and immune-suppressed subtypes, and immune-activated patients were more suitable for anti-PD-1 immunotherapy^33^. The dysfunction of RNA binding proteins (RBPs) in tumorigenesis is important as well. Our prior work constructed RFS-associated RBPs and generated a prognostic signature to help identify patients with high risk and resistance to ARSI^37^. For miRNAs, we evaluated and constructed 15 miRNA-based prognostic signatures in the TCGA-PRAD cohort and validated the findings in the MSKCC cohort, with preferable results^38^. Several genes and miRNAs identified in the abovementioned studies were also used in the current multi-omics study.

The PMOC system contains three subtypes, determined by cluster prediction index analysis and Gap statistics, and the previous three clusters defined by You et al.^13^ and Zhao et al.^15^ were also considered. The clustering results from the ten algorithms are similar and ultimately generated three consensus clusters. The RFS outcomes of the three PMOC subtypes were significantly separated, as were the specific molecular features. The PMOC1 subtype, also called the inflammatory subtype, contains the highest expression levels of immune checkpoint proteins, moderate activated immune-associated pathways, and contains highly activated metabolic pathways of nicotinamide adenine dinucleotide (NAD) biosynthesis cyclooxygenase (COX) arachidonic acid (ARA) metabolism, retinoid metabolism, cortisol biosynthesis, and aldosterone biosynthesis. NAD+ is reported to be one of the major modulators of immune-metabolic circuits^39^, and COX catalyzes the conversion of ARA to prostaglandins, further regulating immunity and lipid metabolism in mammals^40^. The cortisol and aldosterone biosynthesis pathways are also activated in the PMOC1 subtype, which is linked with tumor inflammation^41^.

The PMOC2 tumor-activated subtype demonstrated the worst prognosis, which might be impacted by the activated cell cycle and DNA repair pathways. As early as 2011, the cell cycle control protein cyclin D1 was identified as a component of a 4-gene signature that predicted poor clinical outcome in prostate cancer^42^. Comstock et al. reported that PD-0332991, an orally active CDK inhibitor, can suppress prostate cancer cell proliferation in both hormone-sensitive and castration-resistant contexts^43^. The PMOC2 subtype also contains the most genetic alterations, including mutations of TP53 and APC and copy number amplification of 8q24.21. For the APC mutation, we observed that muted APC leads to the downregulation of its mRNA expression and is associated with accumulated recurrence events. Bruxuoort et al.^44^ also reported the function of APC in a mouse model, and they showed that prostate-specific deletion of Apc can significantly accelerate the tumorigenesis of prostate adenocarcinoma early to 7 months, which might be impacted by Wnt/β-catenin signaling. Re-establishing Apc expression can restore normal development and differentiation of the intestinal crypt epithelium in mice with Apc-deficient intestinal adenomas and adenocarcinomas^45^, which might be a therapeutic target for the clinical treatment of PCa patients with APC mutation and deficiency. Park et al.^46^ reported that FOXA1 is essential for androgen-dependent PCa growth and can be regulated by EZH2 enzymatic inhibitors.

The PMOC3 subtype is likely to be a “balance” subtype, and we observed activated oncogenic signaling pathways, including hypoxia, angiogenesis, epithelial mesenchymal transition, and PI3K/AKT pathways. Moreover, we observed activated proinflammatory (with the inhibitory function of tumorigenesis) pathways, including IL6/JAK/STAT3, IL2/STAT5, Notch and TNF-α signaling. Activated oncogenic and suppression pathways balance the tumor microenvironment and result in a relatively favorable clinical outcome. Additionally, we observed the activation of the androgen response via ARA score and AR activation signature in the PMOC3 subtype, as well as the high response rate of ARSI or bicalutamide treatment. For the regulons and chromatin remodeling networks, we observed the activated regulons of AR, HIF1A, STAT3, and KLF4. Siu et al.^47^ reported that KLF4 expression is directly and transcriptionally upregulated by AR, while AR expression is reciprocally upregulated by KLF4. KLF4 functions as an activator of the androgen receptor through reciprocal feedback. Geng et al.^48^ revealed that glucose-6-phosphate isomerase (GPI) can be repressed by AR in hypoxia and leads to the inhibition of cell growth, while ARSI treatment can restore the expression of GPI and switch the pattern to hypoxia-induced glycolysis. These results indicated that the AR and HIF1α regulons can switch glycose metabolism in the tumor microenvironment.

Finally, we compared our PMOC subtypes with several published PCa molecular subtypes. In the PAM50 subtypes, we revealed a significant difference in the clinical outcome difference of the LumB subtype. Patients with the LumB/PMOC2 subtypes showed a dramatically poorer prognosis than those with the LumB/PMOC1+3 subtype, which might be attributed to the activation of the DNA repair pathway and immunocyte infiltration. Although the LumB subtype is reported to demonstrate poor prognosis and clinical outcomes among PCa subtypes, Zhao et al.^15^ reported that the 10-year actuarial rates corresponding to the biochemical RFS rates were low at 29% for the LumB subtype, 41% for the LumA subtype, and 39% for the basal subtype. However, in the current study, we separated LumB patients into two diverse prognosis subtypes, which is useful for the clinical identification of patients with high risk.

## Conclusion

Taken together, we defined the PMOC system for PCa patients via multi-omics data and consensus results of ten algorithms. The PMOC1 tumor-inflammatory subtype involves the activation of inflammation-associated metabolism pathways and high levels of immune checkpoint proteins. The PMOC2 tumor-activated subtype contains activated cell cycle and DNA repair pathways, a high rate of gene mutation, and 8q24.21 copy number amplification associated with poor prognosis. In the PMOC3 tumor-balanced subtype with the activation of both oncogenic and proinflammatory pathways, the balanced tumor microenvironment was linked with a favorable prognosis, while enrichment of the AR response pathway and AR regulon indicated the suitability of treatment with ARSI. This multi-omics consensus PCa molecular classification can further assist in the precise clinical treatment and development of targeted therapy.

## DECLARATIONS

### Ethics approval and consent to participate

As the data used in this study are publicly available, no ethical approval was required.

### Consent for publication

Not applicable.

### Availability of data and material

The raw data for this study were generated at the corresponding archives. Derived data supporting the findings are available from the corresponding author [LCZ] on reasonable request. Most of the analytic processes in this study are embedded in the R package “*MOVICS*” at https://github.com/xlucpu/MOVICS, which we recently developed for multi-omics integration and visualization.

### Authors’ contributions

Conceptualization, J. M, X. L and C. J; methodology, J. M, X. L, Q. G, and Y. Z; formal analysis, J. M, X. L, Q. G, J. Z and Y. Z; investigation, J. M, X. L, C. J, Y. Z, Z. H, and S. G; writing the original draft, X. L and J. M; visualization, J. M, M. Z; funding acquisition, C. Z, F. Y, and supervision, C. L, and F. Y.

## Acknowledgements

We greatly appreciate the patients and investigators who participated in the corresponding medical project for providing data.

---

**Figure S1. TMB in three prostate cancer subtypes.**

**Figure S2. G-Score of copy number alterations in PMOC1 and PMOC3 subtypes.**

**Figure S3. PVTI expression among tumor and normal samples in pan-cancer.**

**Figure S4. Recognizing the three subtypes by the template genes in four external validation cohorts.**

